# Developmental gene expression differences between humans and mammalian models

**DOI:** 10.1101/747782

**Authors:** Margarida Cardoso-Moreira, Britta Velten, Matthew Mort, David N. Cooper, Wolfgang Huber, Henrik Kaessmann

**Author notes:** Correspondence: M.C.M. or H.K.

## Abstract

Identifying the molecular programs underlying human organ development and how they differ from those in model species will advance our understanding of human health and disease. Developmental gene expression profiles provide a window into the genes underlying organ development as well as a direct means to compare them across species. We use a transcriptomic resource for mammalian organ development to characterize the temporal profiles of human genes associated with distinct disease classes and to determine, for each human gene, the similarity of its spatiotemporal expression with its orthologs in rhesus macaque, mouse, rat and rabbit. We find that half of human genes differ from their mouse orthologs in their temporal trajectories. These include more than 200 disease genes associated with brain, heart and liver disease, for which mouse models should undergo extra scrutiny. We provide a new resource that evaluates for every human gene its suitability to be modeled in different mammalian species.

## Introduction

The genetic programs underlying human organ development are only partially understood, yet they hold the key to understanding organ morphology, physiology and disease [1–6]. Gene expression is a molecular readout of developmental processes and therefore provides a window into the genes and regulatory networks underlying organ development [7, 8]. By densely profiling gene expression throughout organ development, we get one step closer to identifying the genes and molecular processes that underlie organ differentiation, maturation and physiology [9–13]. We also advance our understanding of what happens when these processes are disturbed and lead to disease. Spatiotemporal gene expression profiles provide a wealth of information on human disease genes, which can be leveraged to gain new insights into the etiology and symptomatology of diseases [8,14–16].

Much of the progress made in unraveling the genetic programs responsible for human organ development has come from research in model organisms. Mice and other mammals (e.g., rats and rhesus macaques) are routinely used as models of both normal human development and human disease because it is generally assumed that the genes and regulatory networks underlying development are largely conserved across these species. While this is generally true, there are also critical differences between species during development, which underlie the large diversity of mammalian organ phenotypes [1–6,8]. Identifying the commonalities and differences between the genetic programs underlying organ development in different mammalian species is therefore paramount for assessing the translatability of knowledge obtained from mammalian models to understand human health and disease. Critically, gene expression profiles can be directly compared between species, especially when they are derived from matching cells/organs and developmental stages. Gene expression therefore offers a direct means to evaluate similarities and differences between species in organ developmental programs. While the relationship between gene expression and phenotypes is not linear, identifying when and where gene expression differs between humans and other species will help identify the conditions (i.e., developmental stages, organs, genes) under which model species may not be well suited to model human development and disease.

Here, we take advantage of a developmental gene expression resource [13], which densely covers the development of seven major organs in humans and other mammals, to characterize the spatiotemporal profiles of human disease genes and gain new insights into the symptomatology of diseases. We also determine for each human gene (including disease-associated genes) the similarity of its spatiotemporal expression with that of its orthologs in mouse, rat, rabbit and rhesus macaque. Our analyses and datasets therefore provide a new resource for assessing the suitability of different mammalian species to model the action of individual genes and/or processes in both healthy and pathological human organ development.

## Results

### An expression atlas of human organ development

The resource [13] provides human gene expression time series for seven major organs: brain (forebrain/cerebrum), cerebellum (hindbrain/cerebellum), heart, kidney, liver, ovary and testis (Figure 1A). The time series start at 4 weeks post-conception (wpc), which corresponds to early organogenesis for all organs except the heart (mid-organogenesis), and then cover prenatal development weekly until 20 wpc. The sampling restarts at birth and spans all major developmental milestones, including ageing (Figure 1A; total of 297 RNA-sequencing (RNA-seq) libraries). Matching datasets are available for mouse (316 libraries), rat (350 libraries) and rabbit (315 libraries) until adulthood and for rhesus macaque starting at a late fetal stage (i.e., embryonic (e) day 93, corresponding to 19 wpc human [13]; 154 libraries; Methods).

**Figure 1.**
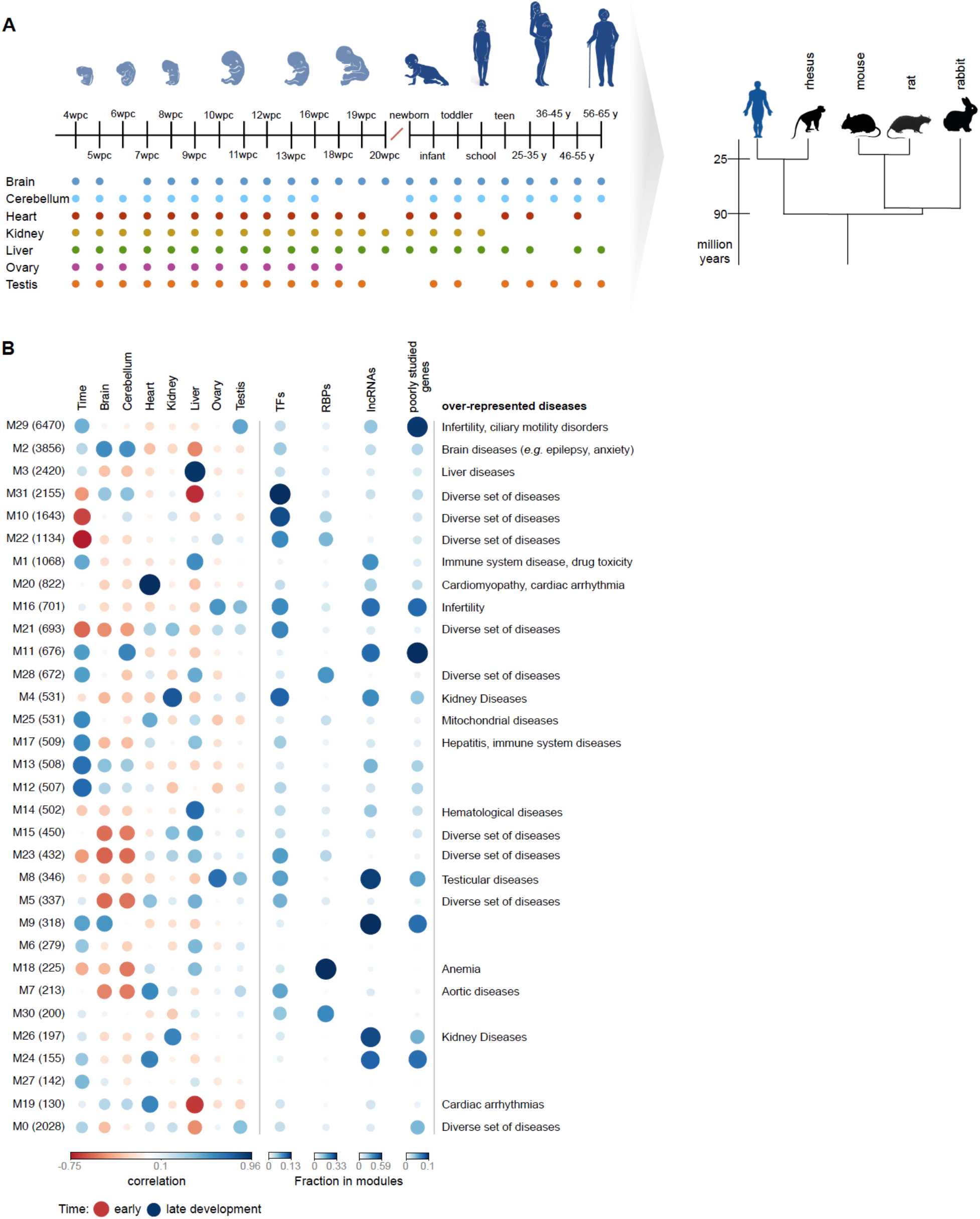
An expression atlas of human organ development. **(A)** Description of the dataset. The dots mark the sampled stages in each organ (median of 2 replicates). **(B)** Modules in the gene co-expression network (number of genes in each module in parentheses), their correlation with organs and developmental time (full developmental profiles in Figure S1A), their fraction of TFs, RBPs, developmentally dynamic lncRNAs and poorly studied protein-coding genes, and examples of overrepresented diseases (FDR < 1%, hypergeometric test; Table S1). The modules are ordered vertically by decreasing number of genes. Module 0 (bottom) includes genes not assigned to any of the other modules.

We used weighted gene co-expression network analysis to identify the main clusters (modules) of highly correlated genes during human organ development (Methods). We then characterized each module according to its developmental profile (Figure 1B; Figure S1A), functional and disease enrichments (Figure 1b; Table S1), and proportion of transcription factors (TFs) [17], RNA-binding proteins (RBPs) [18] and developmentally dynamic long noncoding RNAs (lncRNAs) [19] (Figure 1B). As expected, there is a clear match between the disease enrichments of each module and its organ developmental profile (Figure 1B). For example, module M3 comprises 2,420 genes mainly expressed in the liver and it is associated with a number of liver-related diseases (e.g., fatty liver). Module M20 (822 genes) comprises genes mainly expressed in the heart and is associated with a number of cardiomyopathies. Consistent with previous work [13], we observe that modules associated with higher expression early in development have a significantly higher fraction of TFs than modules associated with higher expression late in development (Pearson’s *ρ*: −0.71, *P*-value *=* 5 x 10^-6^; Figure S1B), a result that is consistent with TFs directing most of organogenesis. The modules identified also provide a wealth of information on poorly characterized genes, that through “guilt-by-association” can be assigned putative functions (Table S2). We identified a strong positive correlation between the fraction of protein-coding genes in a module that are among the least studied in the human genome [20] and the module’s fraction of dynamic lncRNAs (*ρ*: +0.77, *P*-value = 2 x 10^-7^). Modules rich in dynamic lncRNAs and poorly studied protein-coding genes are frequently associated with high expression in the gonads (Figure 1B) but are also found in association with high expression in each of the other organs (e.g., module M9 for brain and module M11 for cerebellum).

The breadth of developmental expression (i.e., the organ- and time-specificity of a gene) informs on gene function, because it is expected to correlate with the spatiotemporal manifestation of phenotypes. TFs, RBPs and members of the seven major signaling pathways all play key roles during development but have distinct spatiotemporal profiles (Figure S2; time- and organ-specificity are strongly correlated [13]). Consistent with previous observations [18], RBPs are generally ubiquitously expressed, with only 6% (100) showing time-and/or organ-specificity (Figure S3). Among these are the developmental regulators LIN28A and LIN28B, which are expressed at the earliest stages across somatic organs; the heart-specific splicing factor RMB20, which has been associated with cardiomyopathy; gonad-specific RBPs predicted to bind to piRNAs; and several members of the ELAV family of neuronal regulators. Signaling genes also tend to be ubiquitously expressed (Figure S2), but they include a higher fraction (19-20%) of time- and/or organ-specific genes than RBPs. As expected, the greatest variation in the breadth of spatiotemporal expression is found among classes of TFs (Figure S2). Myb TFs are mostly ubiquitously expressed (Figure S4), whereas homeobox, POU-homeobox (Figure S5) and forkhead (Figure S6) TFs display high time- and organ-specificity (Figure S2). Although only 16% of zinc finger TFs show spatiotemporal specificity, they constitute ∼1/4 of all time- and/or organ-specific TFs due to their high abundance. Another ∼1/4 corresponds to homeobox TFs, and the remaining half derive from various classes of TFs. Notable among homeobox TFs are the Hox genes, which are critical for pattern specification at the earliest stages of development [21]. In the developmental span examined in our study, Hox genes play an important role during the development of the urogenital system and the early hindbrain (but not cerebrum) (Figure S7).

### Spatiotemporal profiles of disease genes

The breadth of developmental expression can also inform on the etiology and phenotypic manifestation of human diseases. We integrated a dataset of human essential genes [22] with the set of genes associated with inherited disease in the manually curated Human Gene Mutation Database (“disease genes”) [23] to compare the breadth of developmental expression of genes in distinct classes of phenotypic severity (Figure 2A). We found a clear association between expression pleiotropy (i.e., fraction of total samples in which genes are expressed) and the severity of phenotypes (Figure 2B). Essential genes that are not associated with disease are likely enriched for embryonic lethality and are, congruently, the most pleiotropic. Genes that when mutated range from lethality to causing disease (often developmental disorders affecting multiple organs) are less pleiotropic than embryonic lethals but are more pleiotropic than genes only associated with disease (both *P*-value = 2 x 10^-16^, Wilcoxon rank sum test, two-sided; Figure 2B). Finally, non-lethal disease genes are more pleiotropic than genes not associated with deleterious phenotypes (*P*-value = 2 x 10^-5^; Figure 2B). A similar association is obtained when looking independently at organ- and time-specificity (Figure S8A).

**Figure 2.**
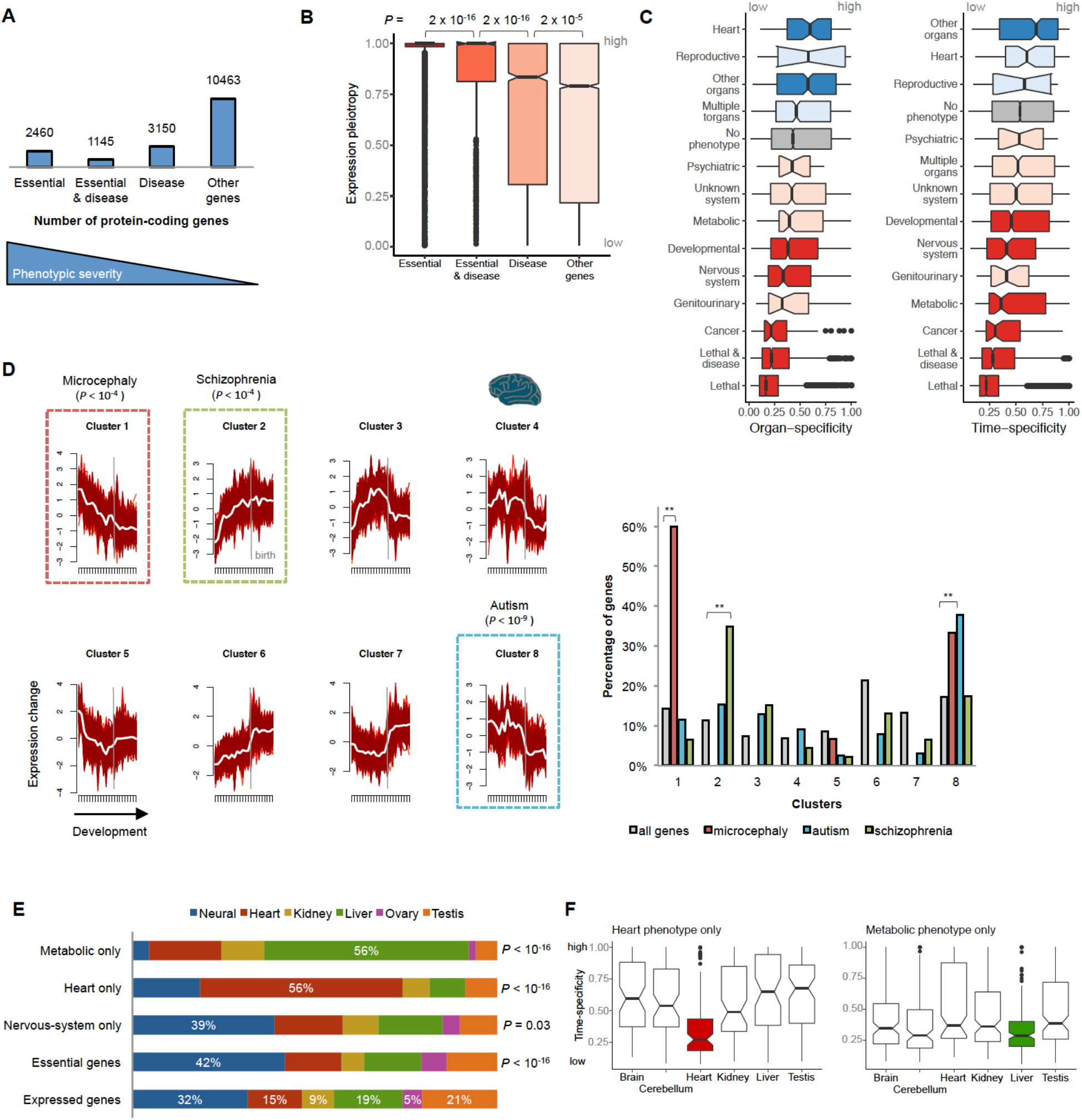
Spatiotemporal profiles of disease genes. **(A)** Number of expressed (RPKM > 1) protein-coding genes in different classes of phenotypic severity. **(B)** Expression pleiotropy of genes in different classes of phenotypic severity (P-values from Wilcoxon rank sum test, two-sided). **(C)** Organ- and time-specificity (median across organs) of genes associated with different classes of diseases. In red are diseases associated with genes with time/organ-specificity lower than non-disease-associated genes and in blue those with higher (darker colors mean that the difference is significant, P ≤ 0.05, Wilcoxon rank sum test, two-sided). **(D)** Genes associated with primary microcephaly (n = 15), autism (n = 164) and schizophrenia (n = 46) are significantly enriched (binomial test) in distinct expression clusters in the brain (on the left are the clusters identified through soft clustering of the brain developmental samples). The genes associated with each disorder are significantly enriched in only one of the 8 clusters (right). **(E)** Organs where genes associated with organ-specific phenotypes show maximum expression. P-values from binomial tests. **(F)** Time-specificity in the different organs of genes with heart- and metabolic-specific phenotypes. In **(B)**, **(C)** and **(F)**, the box plots depict the median ± the 25th and 75th percentiles, with the whiskers at 1.5 times the interquartile range.

Human diseases differ in terms of severity, age of onset and organs affected, all of which should be reflected in the spatiotemporal expression profiles of underlying disease genes. We therefore looked at the time- and organ-specificity of genes associated with different classes of disease [23] (Figure 2C). As expected, the specificity of the spatiotemporal profiles of disease genes differs considerably among disease classes. Genes implicated in developmental disorders, cancer and diseases of the nervous system tend to be ubiquitously expressed (spatially and temporally), whereas genes causing heart and reproductive diseases tend to have more restricted expression (Figure 2C). Further insights were obtained by analysing the temporal trajectories of disease genes within the organs they affect. We used a soft clustering approach to identify the most common expression profiles in each organ and assigned each gene a probability of belonging to each of the clusters (Methods; Table S2). Disease genes are enriched within specific clusters, which are disease and organ-specific. For example, genes associated with heart disease are significantly enriched among genes characterized by a progressive increase in expression throughout heart development, whereas genes associated with metabolic diseases are enriched among genes that exhibit a strong up-regulation in the liver in the first months after birth (Figure S8B-C).

Within the brain, we focused on the temporal trajectories of genes associated with three neurodevelopmental disorders: primary microcephaly, autism and schizophrenia (Methods). Consistent with these disorders having different etiologies and ages of onset, the associated genes are significantly enriched among distinct temporal profiles in the brain (Figure 2D). Genes causing primary microcephaly show their highest expression at the earliest developmental stages followed by a progressive decrease in expression, whereas genes implicated in schizophrenia show the opposite profile: a progressive increase in expression throughout development (Figure 2D). Genes associated with autism are expressed throughout prenatal development and subsequently display a sharp decrease in expression near birth (Figure 2D). The two temporal profiles in the brain that are enriched with microcephaly- and autism-associated genes are also enriched with essential genes (*P*-value < 10^-16^, binomial test).

Most (86%) disease genes that we analyzed are associated with phenotypes in multiple organs, but this still leaves hundreds of genes that affect exclusively one organ. Many of these genes present a puzzle in biomedical research because, as previously noted [24, 25], they are not expressed in an organ-specific manner. Our analysis of developmental transcriptomes further strengthens this puzzle. Genes known to cause organ-specific phenotypes exhibit dynamic temporal profiles in a similar number of organs as genes causing phenotypes across multiple organs (i.e., median of 4 organs for both gene sets; Figure S9A). This raises the question as to why mutations that mostly disrupt the coding-sequences of genes temporally dynamic in multiple organs lead to diseases that are organ-specific. A number of different factors may explain this phenomenon, including alternative splicing (e.g., mutations may affect only organ-specific isoforms) [26], functional redundancy [25], and/or dependency on the characteristics of specific cell types (e.g., protein-misfolding diseases in long-lived neurons). It has also been suggested that pathologies tend to be associated with the organ where the genes display elevated expression [24]. This prompted us to ask where genes associated with organ-specific diseases exhibit their maximum expression during development. We focused on heart, neurodevelopmental, psychiatric, and metabolic diseases (the latter tested in association with the liver). We found a strong association between the organ of maximum expression during development and the organ where the pathology manifests (Figure 2E). Thus, we found that 56% of the genes exclusively associated with heart disease show maximal expression in the heart (vs. 15% for all genes, *P*-value = 2 x 10^-16^, binomial test; Figure 2E), that 56% of the genes with an exclusively metabolic phenotype show maximal expression in the liver (vs. 19% for all genes, *P*-value = 2 x 10^-16^; Figure 2E), and that genes exclusively associated with neurodevelopmental diseases are enriched for maximal expression in the brain (39% vs. 32% for all genes, *P*-value = 0.03; Figure 2D). The duration of gene expression may also help to explain organ-specific pathologies, at least for heart disease. Genes expressed in multiple organs that have heart-specific phenotypes are ubiquitously expressed during heart development but show a significantly higher time-specificity (i.e., shorter expression window) in the other organs (all *P*-value < 10^-4^, Wilcoxon rank sum test, two-sided; Figure 2F). We note, however, that time-specificity does not help to explain metabolic- or neurodevelopmental-specific phenotypes, as we see no difference in the time-specificity of genes in the affected organs versus the others (Figure 2F; Figure S9B). Overall, the association of pathology with level of gene expression, and to a lesser extent with duration of gene expression, suggest that the development of organ-specific pathologies can at least in some cases be explained by differences in the abundance (spatial and/or temporal) of the cell type(s) that express the mutated gene in the different organs.

### Presence/absence expression differences are rare between species

The extensive use of mice, rats and other mammals in biomedical research is predicated upon the assumption of an overall conservation of developmental programs between humans and these species. This assumption has been largely supported by comparative analyses of developmental expression profiles [13] and by comparative analyses of the human and mouse trans-acting regulatory circuitry [27]. However, this broad conservation does not preclude developmental expression differences in individual genes that can profoundly impact the translatability of phenotypes between humans and other species.

We first compared human genes and their orthologs in mouse, rat, rabbit and rhesus macaque in terms of stark differences in spatiotemporal profiles: presence/absence of gene expression in a given organ and large differences in expression pleiotropy across multiple organs. Differences between humans and each of the other species in terms of the presence or absence of gene expression in an organ are rare. In a comparison of human and mouse, only 1-3% of protein-coding genes (177 – 372 genes depending on the organ) are robustly expressed (RPKM ≥ 5) in human but not in mouse (RPKM ≤ 1). These percentages are similar for the comparisons with the other species (i.e., 1-2% of genes robustly expressed in human are not expressed in rat/rabbit/rhesus macaque). Although rare, these differences also include disease genes. For example, among genes robustly expressed in heart in human but not in mouse are 17 genes associated with heart disease. These include *NKX2-6*, which causes conotruncal heart malformations in human [28] that, congruently, are not recapitulated by a mouse knockout [29]. The developmental profile of *NKX2-6* in the human heart is ancestral; the heart expression was lost specifically in rodents, and this is therefore an example of a disease gene that would probably be better studied in the rabbit (Figure 3A). Genes associated with neurological diseases are depleted among the set of genes expressed in the human but not in the mouse brain (12 differ vs. 28 expected, *P*-value = 4 x 10^-4^, binomial test). Among the exceptions is *CHRNA2*, a gene expressed in the human brain starting at birth that has been implicated in epilepsy [30, 31]. Once again, and congruently, this clinical phenotype is not recapitulated in the mouse knockout [29] (Figure 2B).

**Figure 3.**
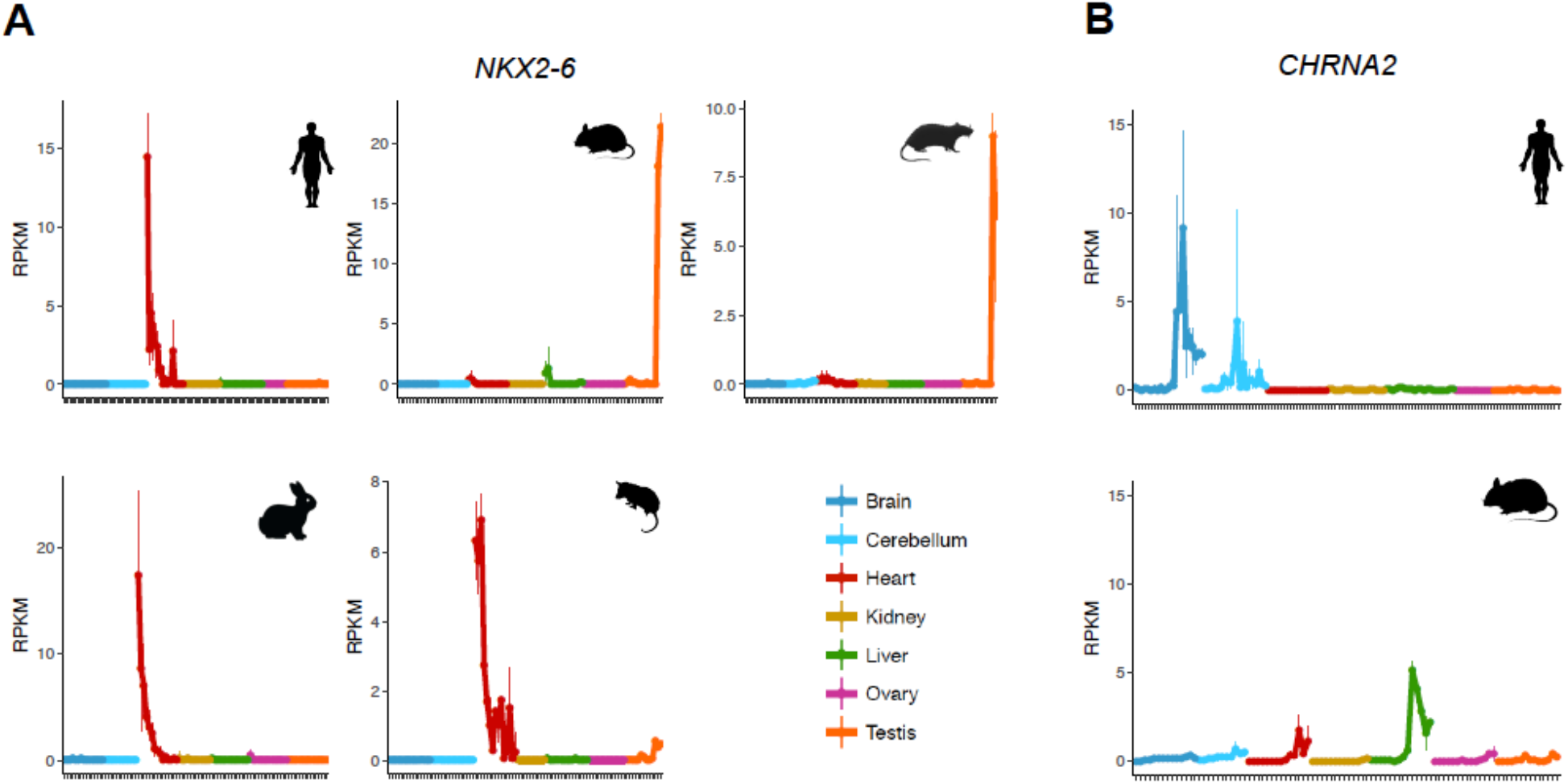
Suitability of the mouse as a model. **(A)** Developmental profile of *NKX2-6* in human, mouse, rat, rabbit and opossum. *NKX2-6* is robustly expressed in the human heart but not in mouse, and the conotruncal heart malformations observed in human are not recapitulated by a mouse knockout. The human heart profile of *NKX2-6* is ancestral as it is similar to the profiles in rabbit and opossum. **(B)** Developmental profile of *CHRNA2* in human and mouse. *CHRNA2* is robustly expressed in the human brain but not in mouse, and the epileptic phenotypes observed in human are not recapitulated by a mouse knockout.

The global breadth of spatiotemporal expression is also very similar between human genes and their orthologs in mouse, rat, rabbit and rhesus macaque. They are highly correlated in terms of their organ-specificity (Pearson’s *r* = 0.85-0.86, all *P*-value < 10^-16^), time-specificity (*r* = 0.68-0.84 for individual organs and 0.83-0.84 for median time-specificity, all *P*-value < 10^-16^) and, therefore, for global expression pleiotropy (*r* = 0.84-0.90, all *P*-value < 10^-16^). There are only 141 genes expressed in at least half the human samples but in fewer than 10% of the mouse samples, and 172 genes with the opposite pattern (Figure S9C). These genes are depleted for essential genes (P-value = 8 x 10^-6^, binomial test) and disease genes (*P*-value = 0.02, binomial test). Overall, differences in the breadth and presence/absence of gene expression between humans and other species are confined to a small set of genes. However, when present, they can translate into relevant phenotypic differences.

### Organ-specific temporal differences are common

It is not uncommon for genes with broad spatiotemporal profiles to evolve new organ developmental trajectories in specific species or lineages [13]. Differences between mammalian species in organ developmental trajectories were first identified using a phylogenetic approach that included distantly related species (i.e., the marsupial opossum which diverged from human ∼160 million years ago [32]) [13]. Therefore, only a restricted set of human genes was evaluated for potential trajectory differences (e.g., 3,980 genes in the brain). Here, we compared the human developmental profiles in each of the organs with their orthologs in mouse, rat, rabbit and rhesus macaque in a pairwise manner (Methods; Tables S3-S8; because of the shorter rhesus macaque’s time series, this analysis was only performed for brain, heart and liver). This allowed us to duplicate or triplicate (depending on the organ) the number of orthologous genes analyzed for differences in their developmental trajectories. Figure 4A shows examples of genes with different developmental trajectories between human and mouse.

**Figure 4.**
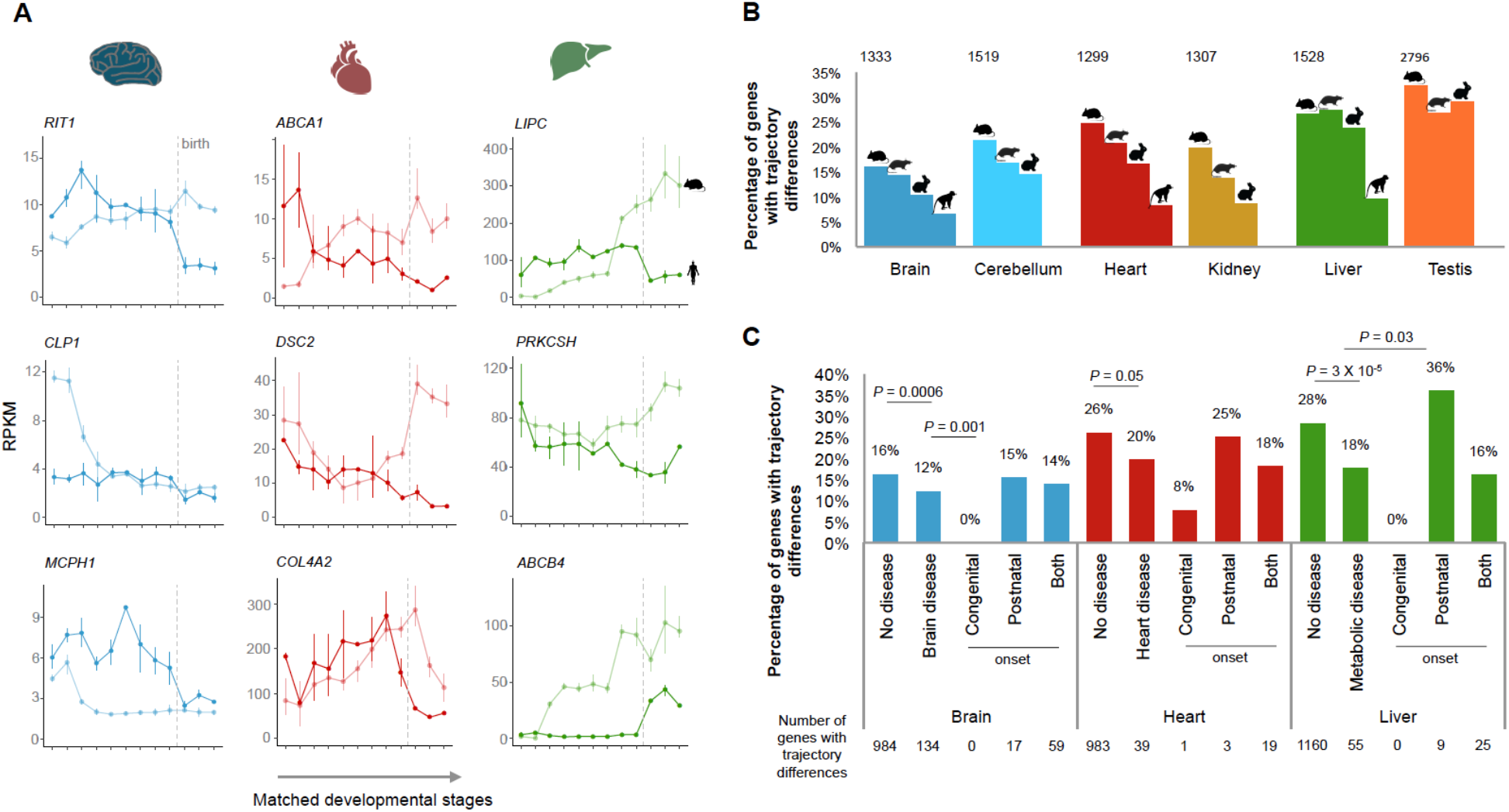
Developmental trajectory differences. **(A)** Examples of human disease genes with different developmental trajectories between human and mouse in the affected organ. **(B)** Percentage of genes in each organ that have different trajectories between human and mouse, rat, rabbit and rhesus. On the top are the number of genes that have a different trajectory between human and mouse. **(C)** Percentage of genes in brain, heart and liver that differ in trajectories between human and mouse. *P*-values for comparisons between disease and non-disease genes are from Fisher’s exact tests and *P*-values for comparisons of disease genes with different ages of onset are from binomial tests.

Consistent with the original study [13], we found differences between the organs in the proportion of genes with trajectories differences between humans and each of the other species (Figure 4B): differences are highest in testis and liver and lowest in brain. There are also expected differences between species: a smaller fraction of genes differs between human and rhesus macaque (diverged ∼29 million years ago) than between human and each of the glires (diverged ∼90 million years ago). However, we also identified a higher proportion of genes that differ between human and mouse than between human and rabbit (despite the same divergence time), a result consistent with the original observation that rodents have evolved a larger number of trajectory differences [13].

Genes with different developmental trajectories between human and mouse are common: 51% of the genes tested differ in at least one organ. Most of these genes (67%) differ in only one organ (25% of genes differ in two organs, and 8% differ in 3 or more), despite on average showing dynamic temporal profiles in 5-6 organs. Genes with differences in developmental trajectories are depleted for TFs (*P*-value = 2 x 10^-5^, Fisher’s exact test, two-sided) and are functionally enriched for protein metabolism (Benjamini-Hochberg corrected *P*-value = 1 x 10^-4^, overrepresentation enrichment analysis). Interestingly, genes with different trajectories in the brain (but not in the other organs) are enriched among a set of genes identified as carrying signs of positive-selection in their coding-sequences across mammalian species [33] (*P*-value = 0.008, Fisher’s exact test).

The genes depicted in Figure 4A are associated with diseases that affect the organ in which human and mouse display different trajectories. For these genes, the disease etiology may not be fully recapitulated by mouse models. The mouse knockouts are still expected to affect the development of the organ associated with the disease, but the cellular and developmental context of the phenotypes in mouse could differ substantially from those in human. It is therefore noteworthy that genes associated with human disease are less likely than non-disease genes to differ in their trajectories between human and mouse (Figure 4C; and between human and the other species; data not shown). Genes causing diseases that affect the brain, heart and liver are significantly depleted for trajectory differences between human and mouse in each of the organs (Figure 4C, *P*-value = 0.0006 for the brain, *P*-value = 0.05 for the heart and *P*-value = 3 x 10^-5^ for the liver, Fisher’s exact test). Nevertheless, that leaves more than 200 disease genes whose developmental profiles may not be fully recapitulated in the mouse (Figure 4C).

We also posed the question as to whether genes underlying diseases with different ages of onset are equally likely to differ between human and mouse. Although the number of disease genes associated with an exclusive congenital or exclusive postnatal onset is low, we found that genes with congenital onsets almost never differ in terms of their developmental trajectories between human and mouse (i.e., only 1 out of 82 genes causing disease in the brain, heart or liver; Figure 4C) whereas genes with postnatal onsets are more likely to show differences (although this difference is only statistically significant for the liver, *P*-value = 0.03, binomial test; Figure 4C). Overall, we suggest that for genes with differences in developmental trajectories (Tables S3-S8), existing mouse models of human diseases should undergo extra scrutiny and the possibility of studying alternative models should be carefully considered.

## Discussion

We integrated datasets of human essential and disease genes with developmental gene expression profiles in order to shed new light on the causes and phenotypic manifestations of human diseases. We found that the breadth of spatiotemporal expression correlates positively with the severity of phenotypes and that it differs considerably among genes associated with different disease classes. We also found that disease-associated genes are enriched within specific developmental modules in the organs affected, and that genes associated with different brain developmental disorders show distinct temporal profiles during brain development. There is therefore a clear association between spatiotemporal profiles and the phenotypic manifestations of diseases.

The analysis of developmental transcriptomes further strengthened the apparent paradox of ubiquitously expressed genes often having organ-specific phenotypes [24, 25]. We could not distinguish genes associated with organ-specific phenotypes from those associated with multiorgan phenotypes based on the breadth of spatiotemporal profiles. However, for genes associated with organ-specific phenotypes, there is a strong association between the organ affected and the organ of maximal expression during development. This association suggests that at least some organ-specific pathologies could be explained by differences between organs in the spatial and temporal abundance of the cells expressing the mutated gene.

Gene expression links genes with their organismal phenotypes and hence offers a direct means to compare both across species. It can, therefore, inform on the likelihood that insights obtained from studies in model species are directly transferable to human. We found that stark changes in gene expression (e.g., presence/absence of expression) are rare between species. However, they do sometimes occur in disease genes, and in these cases, they may explain why for these genes mouse models fail to recapitulate the human phenotypes. Strikingly, we also found that differences between humans and other species in terms of the genes’ temporal trajectories during organ development are common. About half of human genes exhibit a different developmental trajectory from their mouse orthologs in at least one of the organs. In further support of the use of model organisms for disease research, we found that disease genes are less likely to differ than the average gene. Nevertheless, we still identified more than 200 genes known to be causally associated with brain, heart and/or liver disease, that differ in developmental trajectories between human and mouse in the affected organ.

Different reasons, that are not mutually exclusive, can account for the differences in temporal trajectories observed between species. Differences in developmental trajectories can be created by gene expression differences between species in homologous cell types, differences between species in cellular composition, and/or differences between species in the cell types that express orthologous genes. All of these differences can decrease the likelihood that the phenotype associated with a human gene will be fully recapitulated in a model species. However, differences in trajectories created by changes in the identity of the cell types that express an orthologous gene in different species will lead to the greatest phenotypic divergence. Endeavors that seek to clarify the causes of trajectory differences therefore represent a key next step, given that they will identify further genes and processes that are challenging to model in other species. The use of single-cell technologies will greatly aid these efforts [34].

Gene expression is only one of several steps connecting genes to their phenotypes. Similarities and differences in gene expression between species will not always translate into conserved and divergent phenotypes, respectively. This notwithstanding, detailed comparisons of developmental gene expression profiles, as performed here, can substantially help to assess the translatability of the knowledge gathered on individual genes from model species to humans.

## Supporting information

Supplemental Tabes 1-9

## Acknowledgments

We thank S. Anders, R. Arguello, I. Sarropoulos, M. Sanchez Delgado, M. Sepp, T. Studer, Y. E. Zhang and members of the Kaessmann group for discussions. D.N.C and M.M. are in receipt of financial support from Qiagen through a License Agreement with Cardiff University. This research was supported by grants from the European Research Council (615253, OntoTransEvol) and Swiss National Science Foundation (146474) to H.K, and Marie Curie FP7-PEOPLE-2012-IIF to M.C.M. (329902).

## Author Contributions

M.C.M. and H.K. conceived the study. M.C.M. performed the analyses. M.C.M. wrote the manuscript, with input from all authors. B.V. and W.H. contributed to the analyses on trajectory differences. M.M. and D.N.C. contributed to the analyses on human inherited disease.

## Methods

### Resource

From a mammalian resource on organ development [13], we analyzed data from 1,443 strand-specific RNA-seq libraries sequenced to a median depth of 33 million reads: 297 from human, 316 from mouse (outbred strain CD-1 - RjOrl:SWISS), 350 from rat (outbred strain Holtzman SD), 315 from rabbit (outbred New Zealand breed) and 165 from rhesus macaque. The organs, developmental stages and replicates sampled in each species are described in Table S9. The mouse time series started at e10.5 and there were prenatal samples available for each day until birth (i.e., e18.5). There were postnatal samples for 5 stages: P0, P3, P14, P28 and P63. The rat time series started at e11 and there were prenatal samples available for each day until birth (i.e., e20). There were postnatal samples for 6 stages: P0, P3, P7, P14, P42 and P112. The rabbit time series started at e12 and there were 11 prenatal stages available up to and until e27 (gestation lasts ∼ 29-32 days). There were postnatal samples for 4 stages: P0, P14, P84 and P186-P548. Finally, the time series for rhesus macaque started at a late fetal stage (e93) and there were 5 prenatal stages available up to and until e130 (gestation last ∼ 167 days). There were postnatal samples for 8 stages: P0, P23, 5-6 months of age, 1 year, 3 years, 9 years, 14-15 years, and 20-26 years. For mouse, rat and rabbit there were typically 4 replicates (2 males and 2 females) per stage, except for ovary and testis (2 replicates). For human and rhesus macaque, the median number of replicates was 2.

### Gene co-expression networks

We built gene co-expression networks using weighted correlation network analysis (WGCNA 1.61) [35]. We used as input data the read counts after applying the variance stabilizing (VS) transformation implemented in DESeq2 (1.12.4) [36]. Each stage was represented by the median across replicates. In addition to protein-coding genes, we included a set of 5,887 lncRNAs that show significant differential temporal expression in at least one organ and that show multiple signatures for being enriched with functional genes [19]. We only excluded genes that failed to reach an RPKM (reads per kilobase of exon model per million mapped reads) across all stages and organs higher than 1. Using WGCNA we built a signed network (based on the correlation across all stages and organs) using a power of 10 and default parameters. We then correlated the eigengenes for each module with the sample traits (i.e., organ and developmental stage).

We characterized each module in terms of biological processes and disease enrichments (GLAD4U) using the R implementation of WebGestalt (FDR ≤ 0.01; version 0.0.5) [37]. The lists of TFs are from the animalTFDB (version 2.0) [17], the list of RNA-binding proteins are from the work of Gerstberger and colleagues [18], and the lists of genes from the main signaling pathways are from the following Gene Ontology (GO) categories: GO:0016055 (Wnt signaling pathway); GO:0007179 (transforming growth factor beta receptor signaling pathway); GO:0007224 (hedgehog signaling pathway); GO:0007169 (transmembrane receptor protein tyrosine kinase signaling pathway); GO:0030522 (intracellular receptor signaling pathway); GO:0007259 (JAK-STAT cascade); and GO:0007219 (Notch signaling pathway).

### Inherited disease genes

The list of genes associated with human inherited disease was obtained from the manually curated HGMD (PRO 17.1) [23]. We only used genes with disease-causing mutations (DM tag; Table S2). Genes associated with DM mutations were mapped onto the Unified Medical Language System (UMLS), and aggregated into one or more of the following high level disease types: Eye, Nervous system, Reproductive, Cancer, Skin, Heart, Blood, Blood Coagulation, Endocrine, Immune, Digestive, Genitourinary, Metabolic, Ear Nose & Throat, Respiratory, Developmental, Musculoskeletal, and Psychiatric [23]. We also characterized the developmental profiles of 15 genes with dynamic temporal profiles in the brain that are associated with primary microcephaly (out of a set of 16 genes associated with this condition [38]), 171 associated with autism (out of 233 [39]) and 46 associated with schizophrenia (out of 75 [40]; we only considered loci where at most two genes were associated with the causative variant). There were only 7 genes with dynamic temporal profiles in the brain associated with both autism and schizophrenia. The list of human essential genes was obtained from the work of Bartha and colleagues [22].

The time- and organ-specificity indexes were based on the Tau metric of tissue-specificity [41] and were retrieved from the developmental resource [13]. Both indexes range from 0 (broad expression) to 1 (restricted expression). The pleiotropy index is to the number of samples where a gene is expressed (RPKM > 1) over the total number of samples.

The most common temporal profiles in each organ were identified using the soft-clustering approach (c-means) implemented in the R package mFuzz (2.32.0) [42, 43]. The clustering was restricted to genes previously identified as showing significant temporal differential expression in each organ (i.e., developmentally dynamic genes) [13]. We used as input the VS-transformed counts. The number of clusters was set to 6-8 depending on the organ.

### Comparing developmental trajectories

For each organ, we compared the developmental trajectories of orthologous genes previously identified as showing significant temporal differential expression [13]. We used as input the VS-transformed counts (median across replicates) for matching stages between human and each of the other species. The developmental stage correspondences across species were retrieved from the developmental resource [13]. We used GPClust [44–46], which clusters time-series using Gaussian processes, to cluster the combined data for human and each of the other species. We set the noise variance (k2.variance.fix) to 0.7 and let GPClust infer the number of clusters. For each gene, GPClust assigned the probability of it belonging to each of the clusters. Therefore, for each gene we obtained a vector of probabilities that could be directly compared between pairs of 1:1 orthologs. We calculated the probability that pairs of orthologs were in the same cluster and used an FDR cut off of 5% to identify the genes that differed in trajectory between human and each of the other species. In Tables S3-S8, we provide for each organ and species the *P*-values (adjusted for multiple testing using the Benjamini-Hochberg procedure [47]) for the null hypothesis that orthologs have the same trajectory, and their classification as ‘same’ or ‘different’ based on an FDR of 5%.

General statistics and plots. Statistical analyses and plots were done in R (3.3.2) [48]. Plots were created using the R packages ggplot2 (2.2.1) [49], gridExtra (2.2.1) [50], reshape2 (1.4.2) [51], plyr (1.8.4) [52], and factoextra (1.0.4) [53].

## Supplementary figure legends

**Figure S1.**
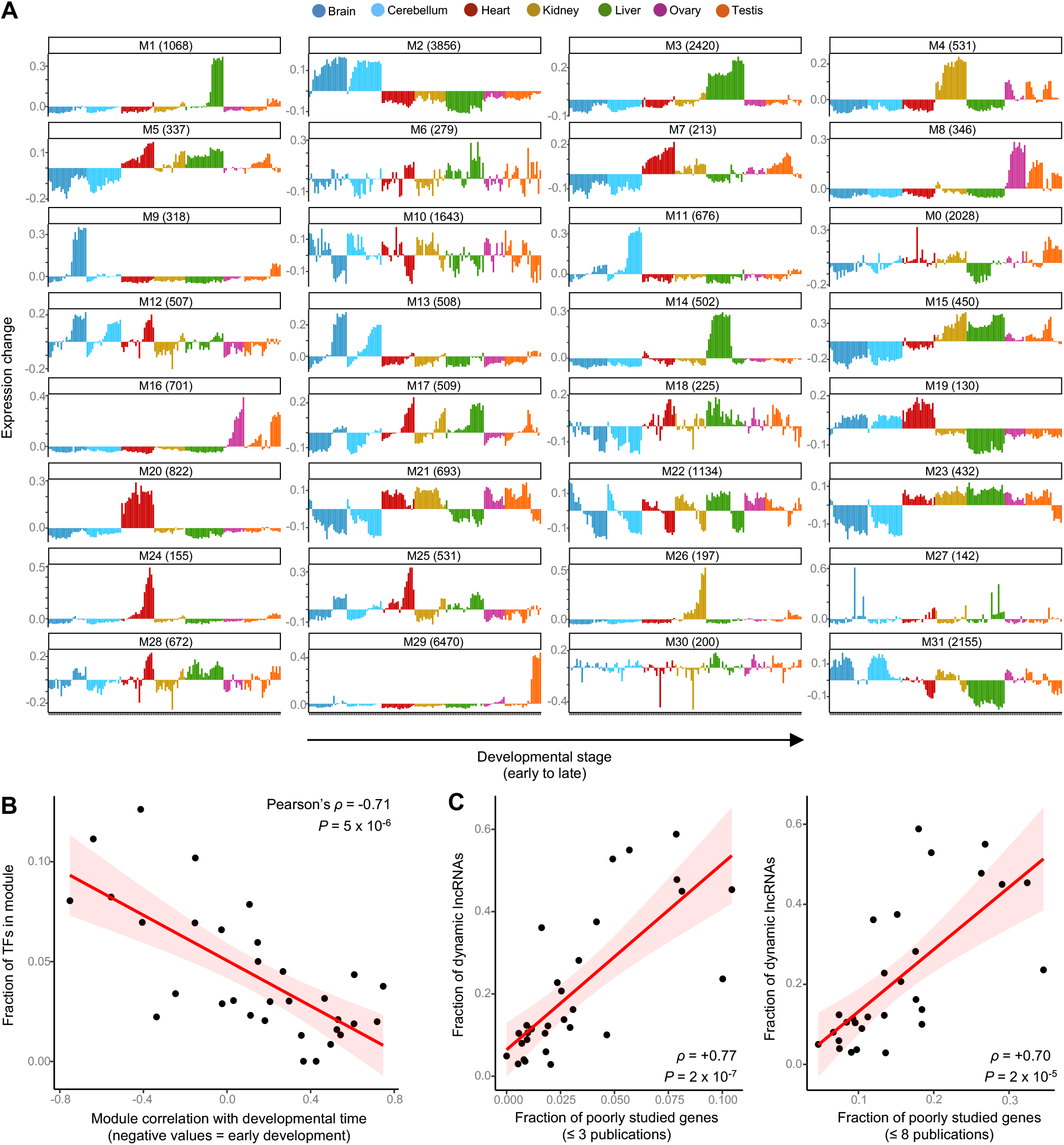
Human weighted gene co-expression network. **(A)** Organ developmental profiles for each module; shown is the module’s eigengene. **(B)** Modules with a high fraction of TFs are associated with expression in early development whereas modules with a low fraction of TFs are associated with expression in late development. Shaded area corresponds to the 95% confidence interval. **(C)** There is a strong positive correlation between the fraction of developmentally dynamic lncRNAs in a module and the fraction of poorly studied protein-coding genes. Poorly studied genes are those with 3 or fewer publications (left) or those with 8 or fewer publications (right). Data on the number of publications are from Stoeger and colleagues. Shaded area corresponds to the 95% confidence interval.

**Figure S2.**
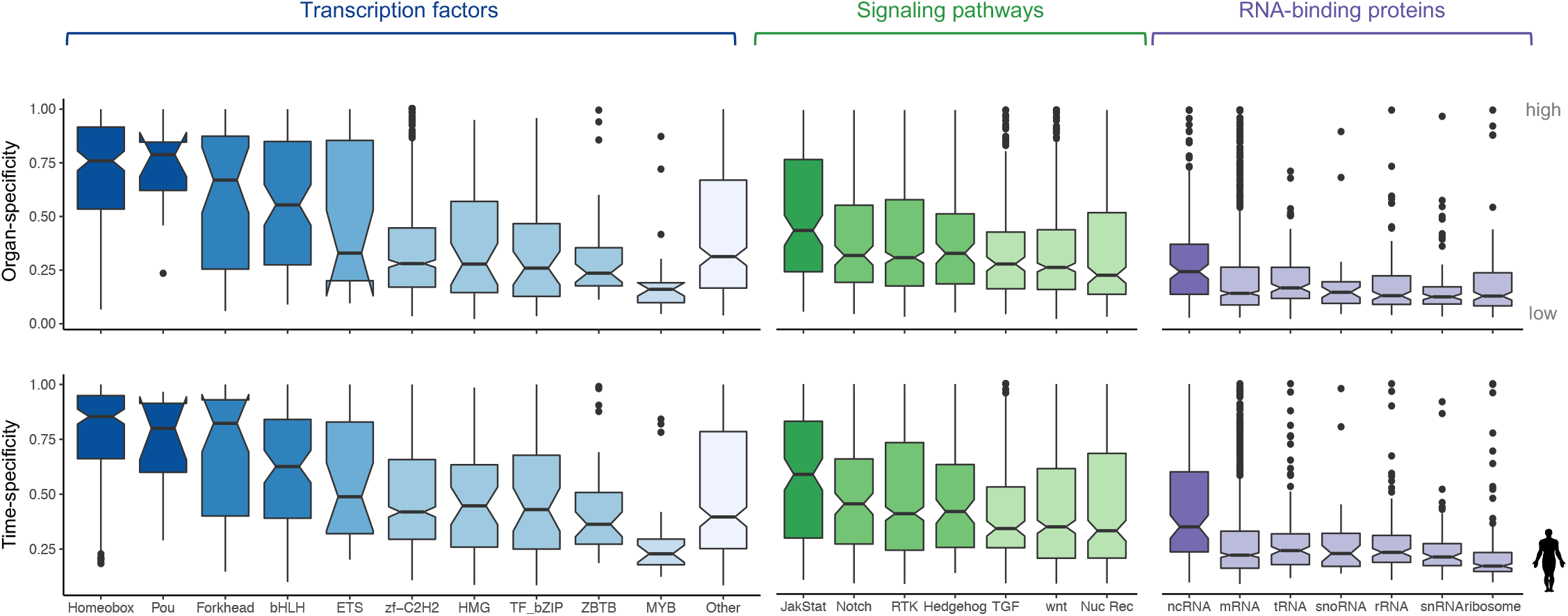
Breadth of developmental expression of key groups of developmental genes. Time and organ-specificity of selected sets of TFs, signaling genes and RBPs. Both indexes range from 0 (broad expression) to 1 (restricted expression) (Methods). The boxplots depict the median ± 25th and 75th percentiles, whiskers at 1.5 times the interquartile range.

**Figure S3.**
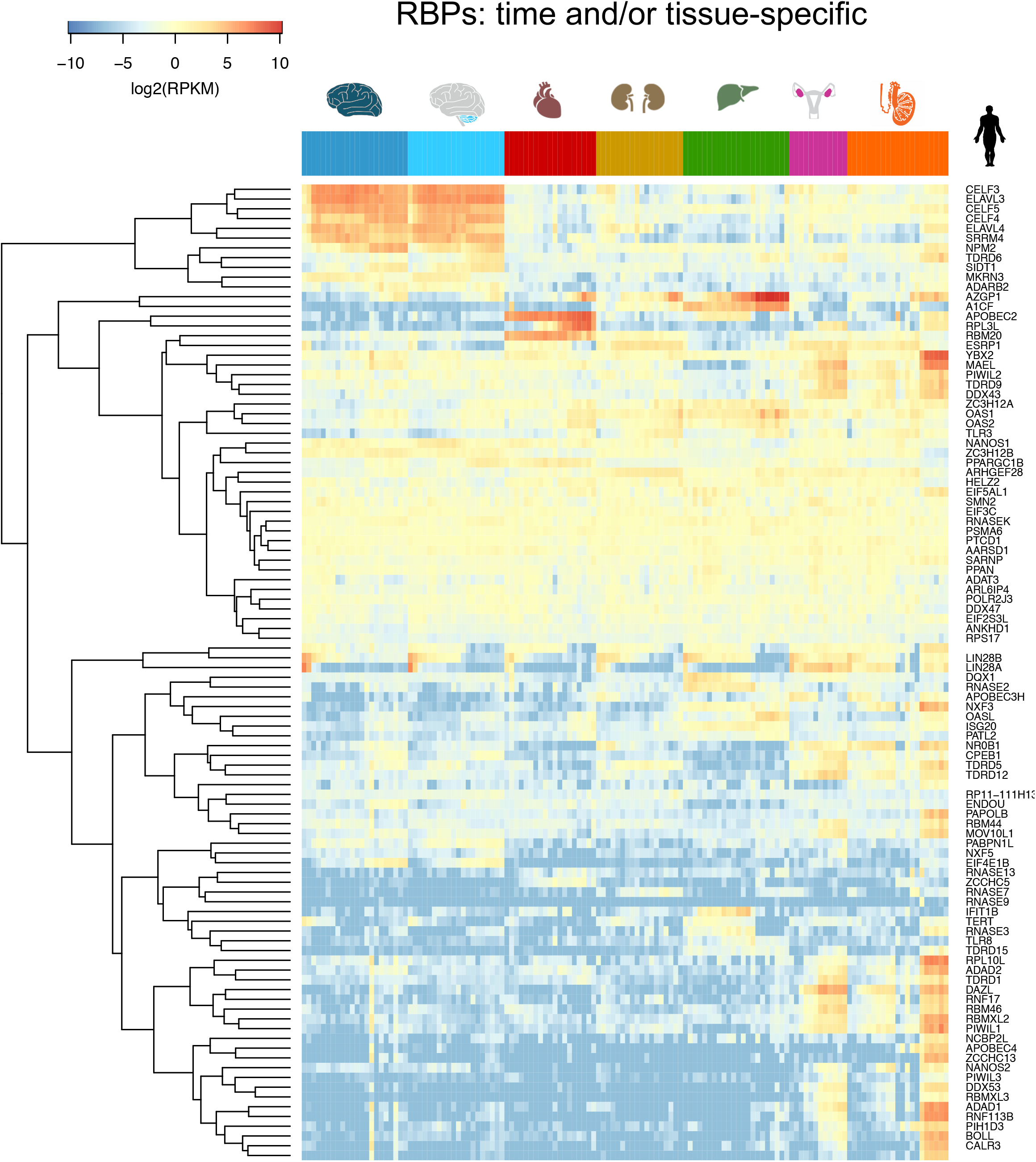
Spatiotemporal profile of time and/or organ-specific RBPs. (organ- and/or median time-specificity ≥ 0.8). In each organ, the samples are ordered from early to late development.

**Figure S4.**
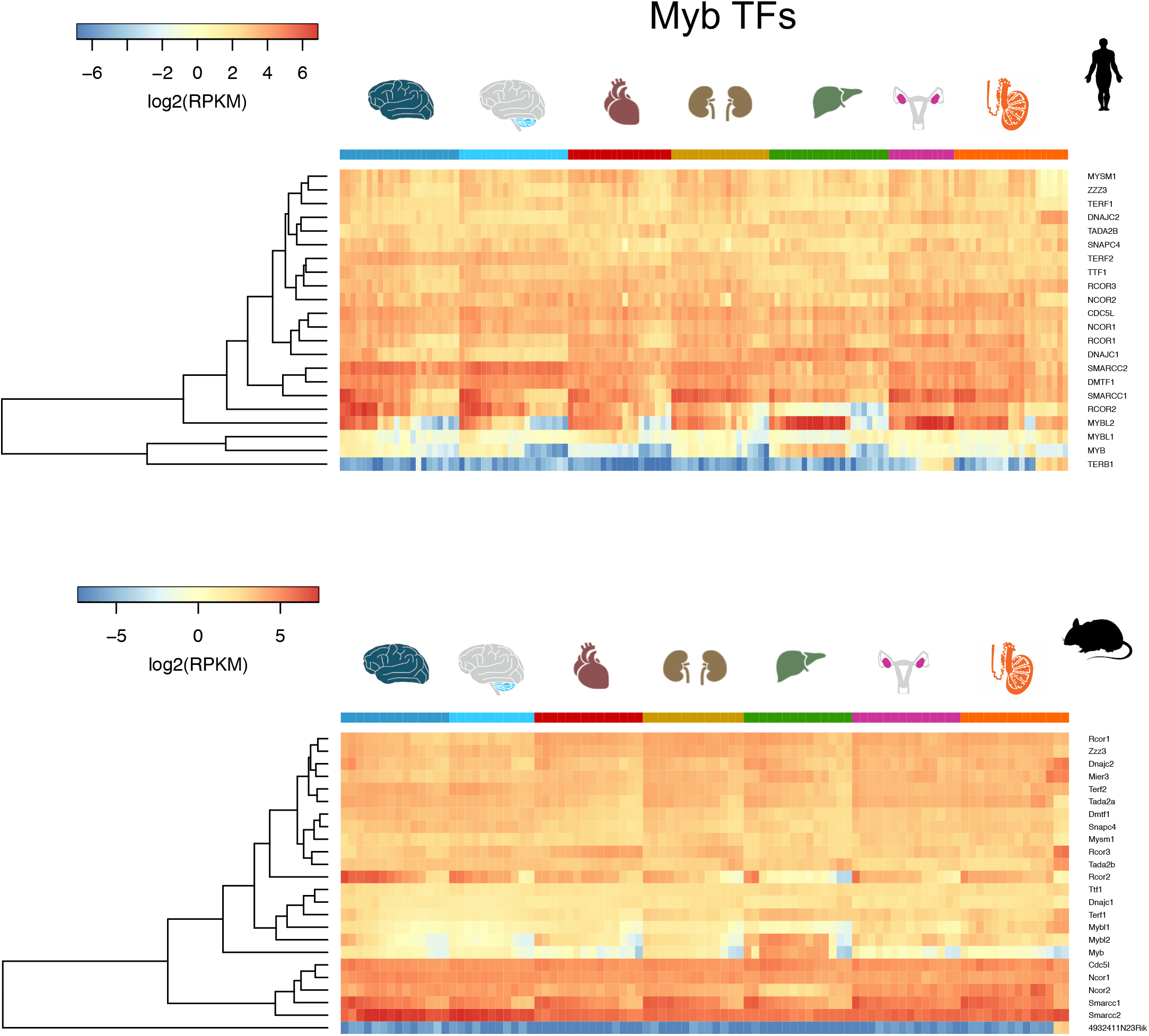
Spatiotemporal profiles of TFs with a Myb DNA binding domain. In each organ, the samples are ordered from early to late development.

**Figure S5.**
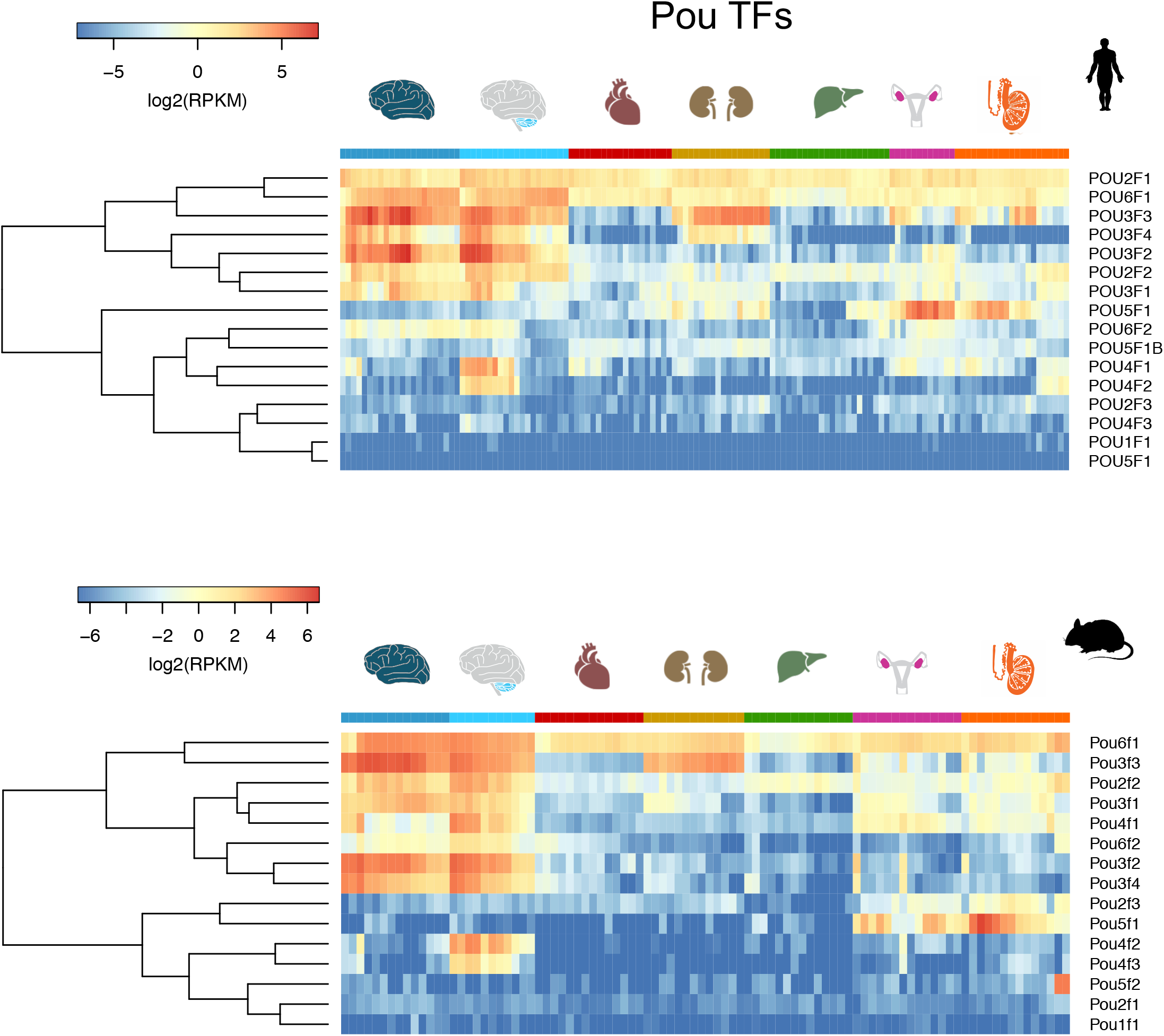
Spatiotemporal profiles of TFs with a POU domain. In each organ, the samples are ordered from early to late development.

**Figure S6.**
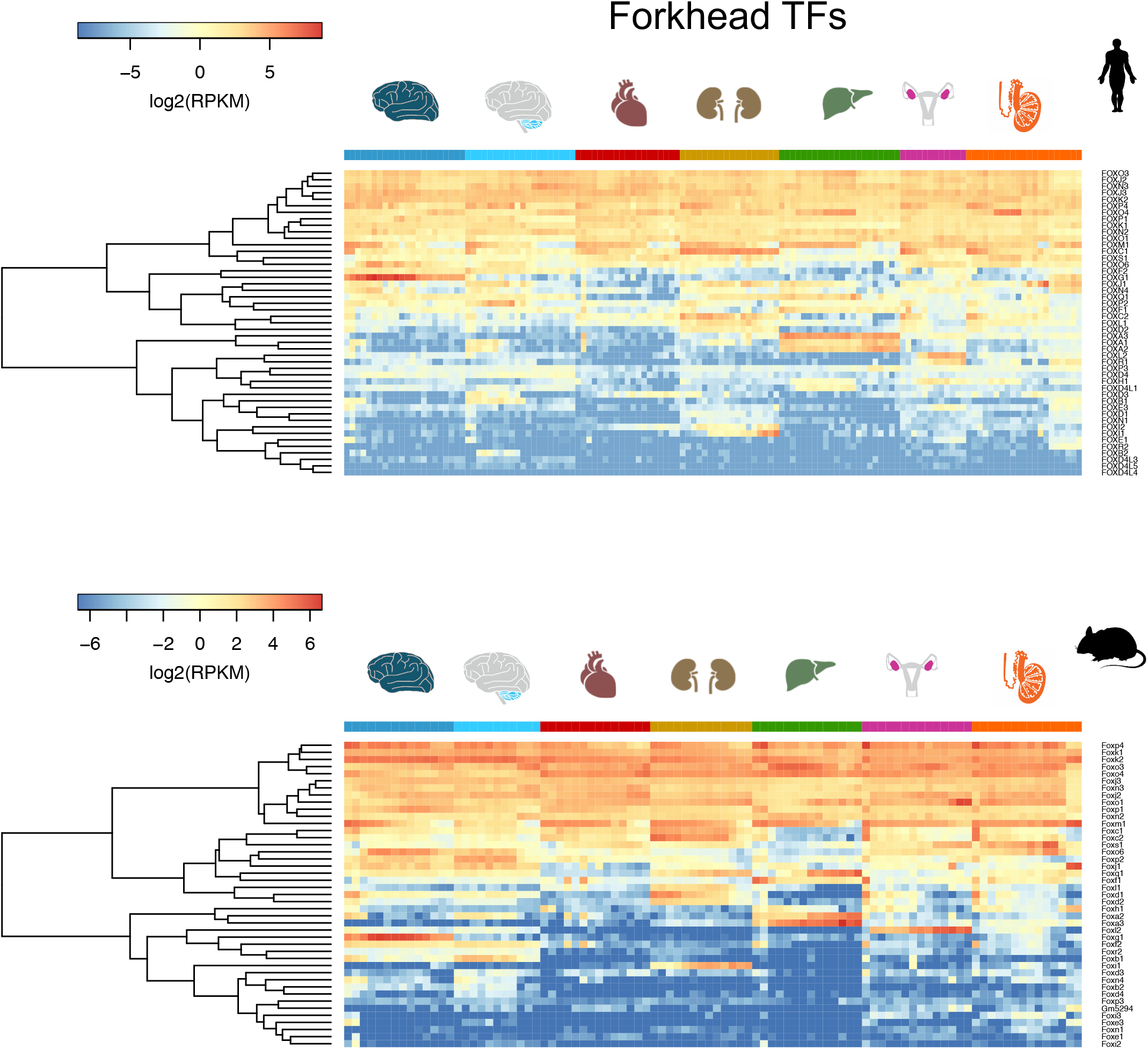
Spatiotemporal profiles of TFs with a Forkhead domain. In each organ, the samples are ordered from early to late development.

**Figure S7.**
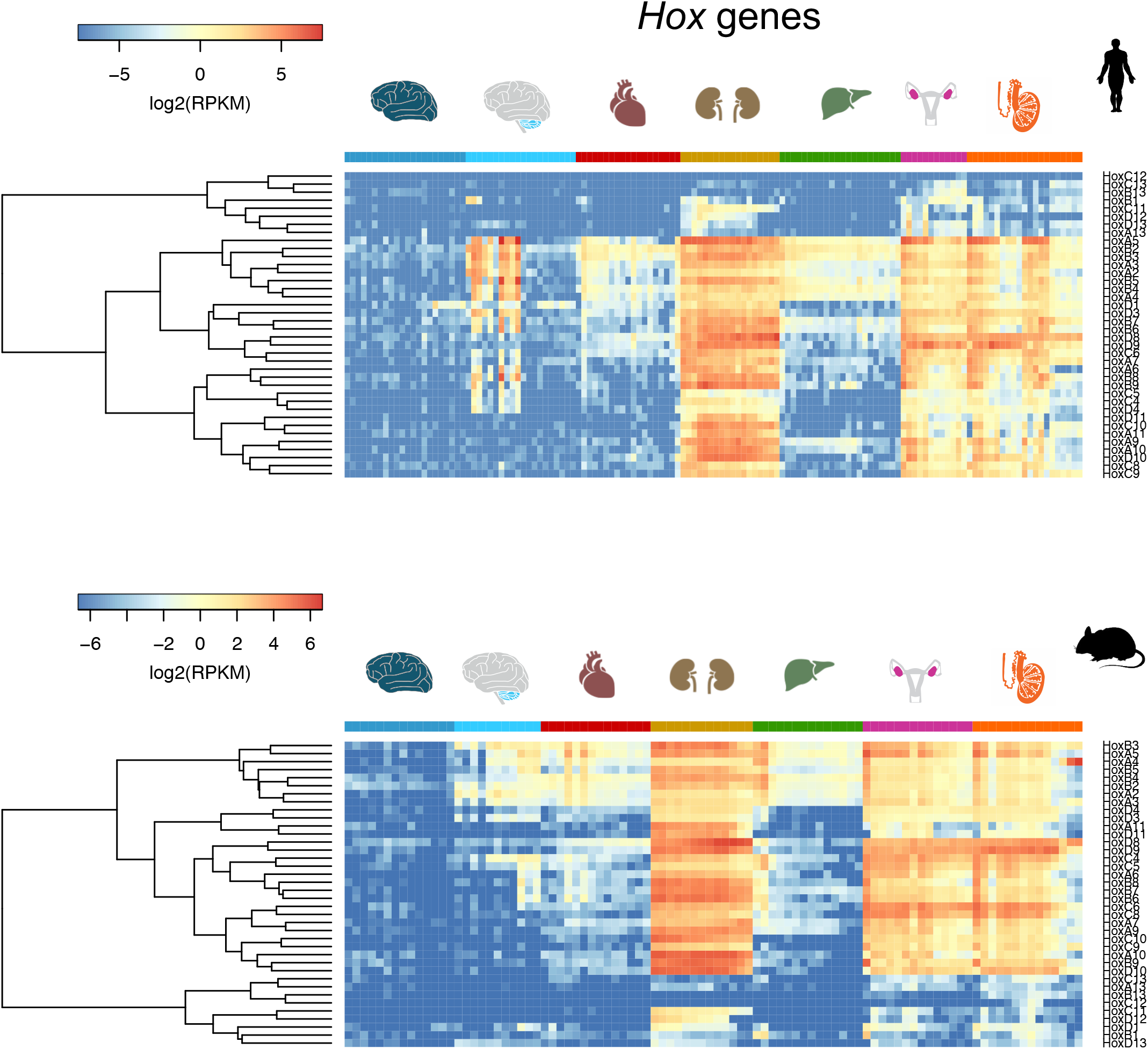
Spatiotemporal profiles of Hox genes. In each organ, the samples are ordered from early to late development.

**Figure S8.**
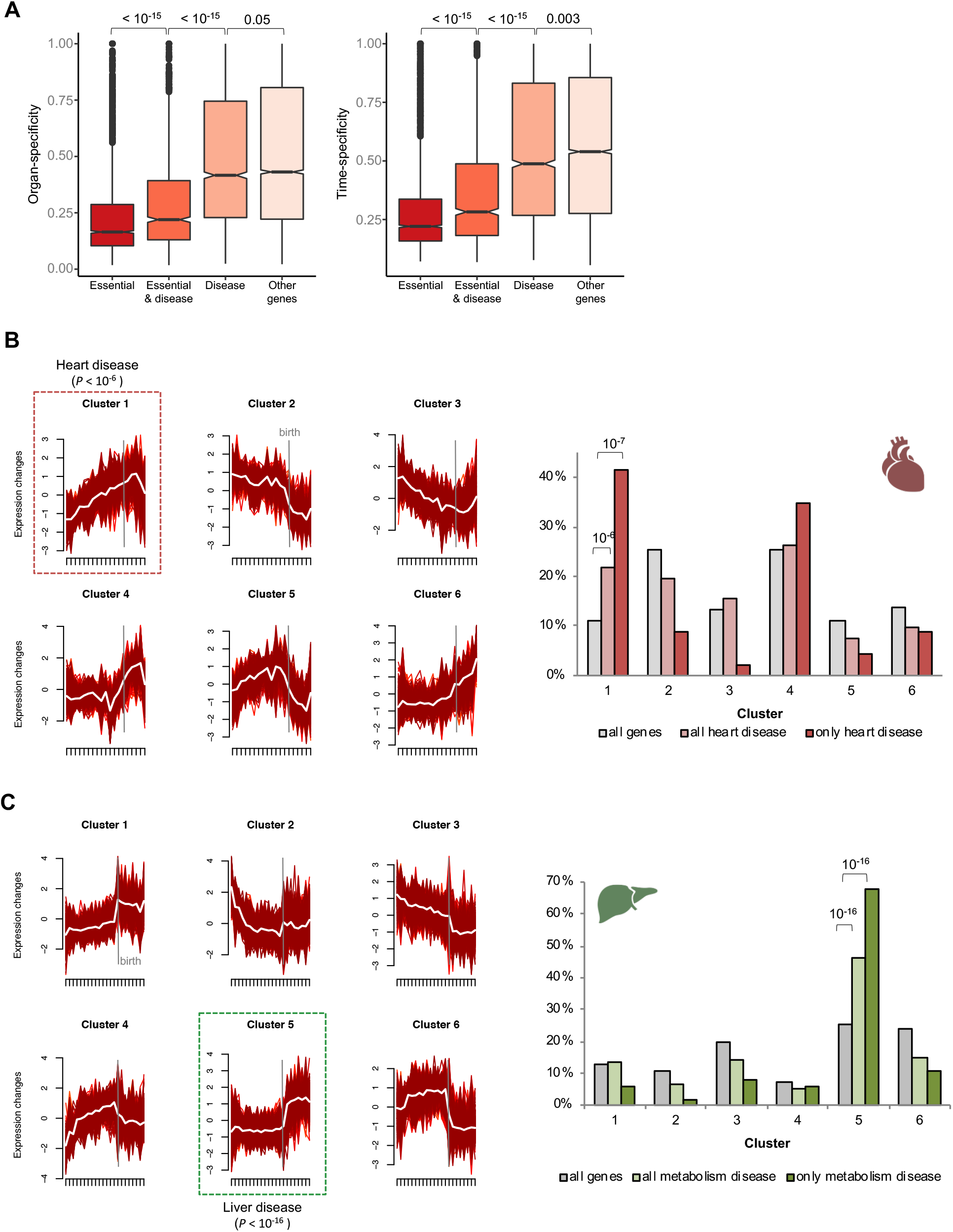
Spatiotemporal profiles of disease genes. **(A)** Organ- and time-specificity (median across organs) for genes in different classes of phenotypic severity (P-values from Wilcoxon rank sum test, two-sided). The boxplots depict the median ± 25th and 75th percentiles, whiskers at 1.5 times the interquartile range. **(B)** Distribution of genes associated with heart disease among the 6 heart clusters. Cluster 1 is enriched for heart disease-associated genes both when using all genes associated with a heart phenotype (n = 230) and when restricting the set to those exclusively associated with the heart (n = 46) (P-values from binomial tests). **(C)** Distribution of genes associated with metabolic diseases among the 6 liver clusters. Cluster 5 is enriched for metabolic disease-associated genes both when using all genes associated with a metabolic phenotype (n = 379) and when restricting the set to those exclusively associated with metabolism (n = 103) (*P*-values from binomial tests).

**Figure S9.**
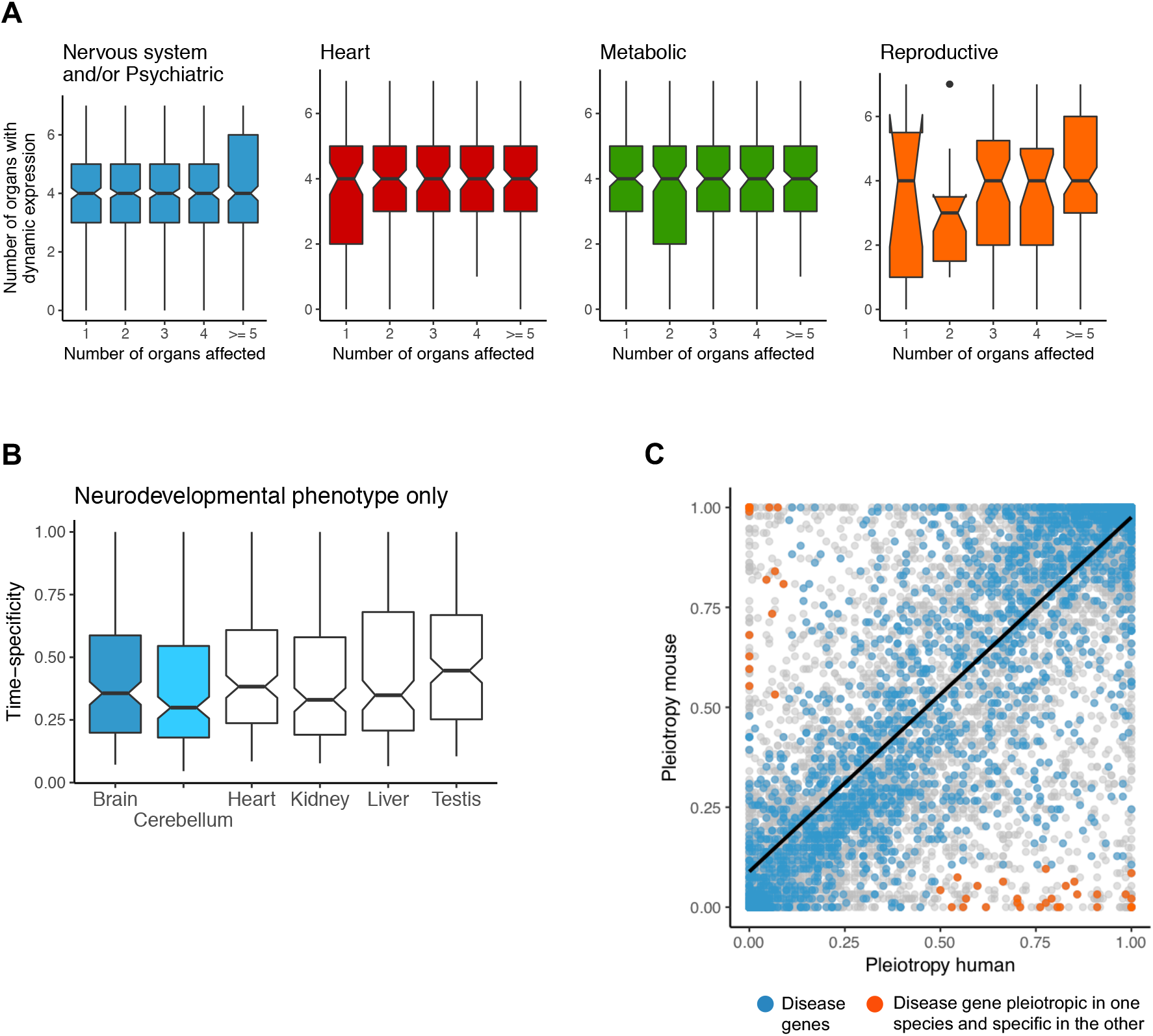
Spatiotemporal profiles of disease genes. **(A)** Number of organs where genes have dynamic temporal profiles as a function of the number of organs where they are known to cause disease. **(B)** Time-specificity in different organs for genes associated exclusively with neurodevelopmental phenotypes. **(C)** Relationship between human and mouse expression pleiotropy. The blue dots denote disease-associated genes and the orange dots denote disease-associated genes expressed in at least 50% of the samples in one species but in less than 10% of the samples in the other. In **(A)** and **(B)** the boxplots depict the median ± 25th and 75th percentiles, whiskers at 1.5 times the interquartile range.

## Supplementary table legends

**Table S1.** Top 5 biological processes and disease enrichments (FDR < 1%, hypergeometric test) for each of the 32 modules in the gene co-expression network.

**Table S2.** Lists of human genes, the modules to which they belong in the global gene co-expression network, the clusters to which they were assigned in each organ (soft clustering), their associations with disease (‘DM’ means disease-causing), the number of organs where they show dynamic temporal profiles, the organ of maximal expression during development, their organ- and time-specificity, and their global expression pleiotropy.

**Table S3**. Comparison of brain temporal trajectories between human genes and their orthologs in mouse, rat, rabbit, and rhesus macaque. Trajectories were called different when the adjusted probability of the orthologs being in the same cluster is ≤ 0.05. Only genes with dynamic temporal profiles in the brain of humans and at least one of the other species were tested for trajectory differences.

**Table S4.** Comparison of cerebellum temporal trajectories between human genes and their orthologs in mouse, rat, and rabbit. Trajectories were called different when the adjusted probability of the orthologs being in the same cluster is ≤ 0.05. Only genes with dynamic temporal profiles in the cerebellum of humans and at least one of the other species were tested for trajectory differences.

**Table S5.** Comparison of heart temporal trajectories between human genes and their orthologs in mouse, rat, rabbit, and rhesus macaque. Trajectories were called different when the adjusted probability of the orthologs being in the same cluster is ≤ 0.05. Only genes with dynamic temporal profiles in the heart of humans and at least one of the other species were tested for trajectory differences.

**Table S6.** Comparison of kidney temporal trajectories between human genes and their orthologs in mouse, rat, and rabbit. Trajectories were called different when the adjusted probability of the orthologs being in the same cluster is ≤ 0.05. Only genes with dynamic temporal profiles in the kidney of humans and at least one of the other species were tested for trajectory differences.

**Table S7.** Comparison of liver temporal trajectories between human genes and their orthologs in mouse, rat, rabbit, and rhesus macaque. Trajectories were called different when the adjusted probability of the orthologs being in the same cluster is ≤ 0.05. Only genes with dynamic temporal profiles in the liver of humans and at least one of the other species were tested for trajectory differences.

**Table S8.** Comparison of testis temporal trajectories between human genes and their orthologs in mouse, rat, and rabbit. Trajectories were called different when the adjusted probability of the orthologs being in the same cluster is ≤ 0.05. Only genes with dynamic temporal profiles in the testis of humans and at least one of the other species were tested for trajectory differences.

**Table S9.** Organs, developmental stages, and number of replicates sampled in each species.

